# Early neutrophilia marked by aerobic glycolysis sustains host metabolism and delays cancer cachexia

**DOI:** 10.1101/2021.08.11.455851

**Authors:** Michele Petruzzelli, Miriam Ferrer, Martijn Schuijs, Zoe Hall, David Perera, Shwethaa Raghunathan, Michele Vacca, Edoardo Gaude, Michael J. Lukey, Duncan I. Jodrell, Christian Frezza, Erwin F. Wagner, Tobias Janowitz, Timotheus Y.F. Halim, Ashok R. Venkitaraman

## Abstract

An elevated neutrophil-to-lymphocyte ratio negatively predicts the outcome of patients with cancer and is associated with cachexia, the terminal wasting syndrome. Here, we show using murine model systems of colorectal and pancreatic cancer that neutrophilia in the circulation and multiple organs, accompanied by extramedullary hematopoiesis, is an early event during cancer progression. Transcriptomic and metabolic assessment reveals that neutrophils in tumor-bearing animals utilize aerobic glycolysis, alike to cancer cells. Although pharmacological inhibition of aerobic glycolysis slows down tumor growth in C26 tumor-bearing mice, it precipitates cachexia, thereby shortening overall survival. This negative effect may be explained by our observation that acute depletion of neutrophils in pre-cachectic mice impairs systemic glucose homeostasis secondary to altered hepatic lipid processing. Thus, changes in neutrophil number, distribution and metabolism play an adaptive role in host metabolic homeostasis during cancer progression. Our findings provide insight into early events during cancer progression to cachexia, with implications for therapy.

## INTRODUCTION

Clinical studies have identified consequences of inflammation, such as changes in the neutrophil-to-lymphocyte ratio (NLR) as an early biomarker of cancer progression^1,2^. Sustained systemic inflammation persists throughout cancer progression and mediates many tumor-host interactions. The cytokines and chemokines of the inflammatory responses in tumors not only contribute to the organization of the tumor microenvironment^3–6^ but also systemically regulate non-tumoral host tissues and their metabolism^7,8^. In particular, systemic inflammation has been implicated in cancer cachexia, which is a common late manifestation of cancer progression defined by wasting of lean body mass and significant weight loss, and is associated with poor prognosis^9^. In patients with cancer and in murine models, elevated interleukin-6 (IL-6) has been causally linked to cachexia and altered macronutrient processing. Mechanistically, IL-6 induces ketogenic impairment due to reprogramming of the liver^7^ and browning of adipose tissue^8^. Also, IL-6 impairs intestinal barrier function in mouse models of cachexia, leading to tranlocation of microbial compounds^10^, which can promote systemic inflammation^11^. Evidence of this reprogramming is already detectable in the pre-cachectic stage. Despite these clear implications of inflammation and the resulting immunological changes in the host, a potential contribution of the immune system to the host biology during cancer progression and ultimately onset of cancer cachexia has not been systematically investigated. Efforts to understand the cellular and systemic processes that precede and drive the development of cancer progression to cachexia, as well as the biology of the pre-cachectic state, are important because overt skeletal muscle catabolism marks an irreversible endpoint of the disease^12,13^.

Here, we characterize the temporal sequence of quantitative and qualitative changes in neutrophils during cancer progression and their role in the pathophysiology of cachexia. Using *in vivo* models that recapitulate the progression of cancer in humans, we report an early and robust increase in circulating neutrophils and widespread neutrophilia in many solid organs that precedes end-stage cachexia. These neutrophils display an enhanced aerobic glycolytic profile that both cancer and immune cells are metabolically dependent on^14,15^, highlighting the need to indentify systemic consequences when targeting this pathway therapeutically. Pharmacological inhibition of aerobic glycolysis resulted in reduced tumor growth, but shortened survival due to early onset cachexia. Targeted depletion of neutrophils revealed a role of inflammatory neutrophilia in lipolysis, lipid accumulation in the liver, and systemic glucose homeostasis. Taken together, our results position neutrophil metabolism at the interface of the metabolic interactions between cancer and host.

## RESULTS

### Widespread neutrophilia is an early systemic manifestation of cancer progression

An increase in the NLR is a well-recognized prognostic marker in patients with advanced cancer^16^ and is associated with cachexia^17^. We utilized the non-metastatic subcutaneous colorectal cancer-derived C26 model, a well-validated murine model of epithelial cancer-induced cachexia associated with inflammation, to dynamically characterize alterations in neutrophils at different stages of disease. Mice were sacrificed at three distinct time points of cancer progression (Fig 1A): an **early** time point (9–10 days post-injection of C26 cancer cells, when mice had small tumors), the **pre-cachectic** stage (15–16 days post-injection, when mice were weight stable with normal food intake), and during **cachexia** (≥21 days post-injection, when mice displayed weight loss and reduced food intake) (Fig 1B, C and Fig EV1A, B). Cachexia was defined by a >10% decrease in total body weight. Splenomegaly developed progressively from the early time point to the cachectic stage (Fig 1D). Cachectic mice exhibited pronounced loss of gonadal white adipose tissue (gWAT) and muscle mass wasting, while pre-cachectic mice showed loss of gWAT but no muscle wasting; gonadal WAT and skeletal mass were unchanged at the early time point (Fig 1E, F). We subseqently performed detailed immunophenotyping of tumours, blood and peripheral organs by flow cytometry to investigate changes in immune profile that precede the onset of cachexia. We detected an increase of blood neutrophils starting at the pre-cachectic stage (Fig 1G). Analysis of tissue neutrophils showed that in the lung and liver of C26 tumor-bearing mice, neutrophil numbers were already increased at the early time point (Fig 1H, I).

**Fig1.**
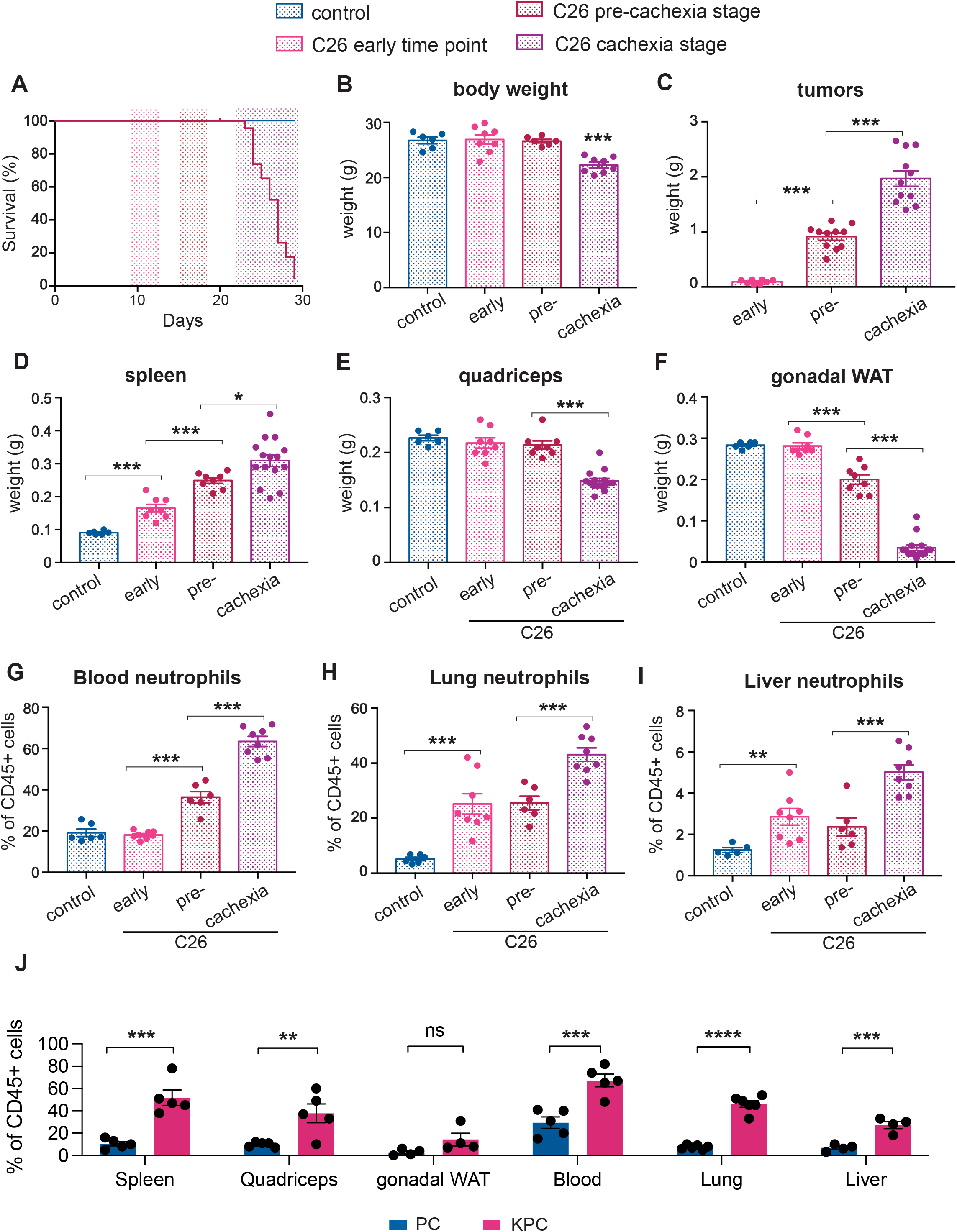
Immune changes during cancer progression in mouse models of cachexia. (A) Mice were injected with C26 colorectal cancer cells and sacrificed at 3 distinct time points: an early time point (9–10 days post-tumor inoculation), the pre-cachectic stage (15–16 days post-injection), and when cachexia occurred (≥21 days post-injection); (B-F) Body (B), tumor (C), spleen (D), quadriceps (E), and gonadal white adipose tissue (gWAT) (F) weights of mice sacrificed at the time points defined in Figure 1A; (G-I) Quantification of neutrophils (displayed as % of neutrophils out of all CD45+ cells), by flow cytometry in the blood (G), lung (H), and liver (I) of mice sacrificed at the time points defined in Figure 1A; (J) Quantification of neutrophils (displayed as % of neutrophils out of all CD45+ cells) in the spleen, quadriceps, gWAT, blood, lung and liver of pre-cachectic KPC mice and PC controls.

We also characterized systemic immune changes at the pre-cachectic stage in a genetically engineered, autochthonous murine model of pancreatic ductal adenocarcinoma (PDAC) that recapitulates human pathology, including the development of cachexia^6^. LSL-KrasG12D/+; LSL-Trp53R172H/+; Pdx1-Cre mutation-bearing (KPC) mice were sacrificed at a timepoint representing pre-cachexia, when tumors of 5-10 mm crossectional diameter were confirmed by ultrasound scanning (Fig EV1C), but no bodyweight or muscle loss was evident when compared to control age- and sex-matched Trp53R172H/+; Pdx1-Cre (PC) littermates (Fig EV1D, E). Despite the absence of weight loss or sarcopenia, we observed a substantial systemic increase in neutrophils in all analyzed tissues of pre-cachectic tumour bearing mice, compared to non-tumour bearing PC controls (Figure 1J). Other immune cell-types were not globally affected; however, eosinophils and macrophages were increased in tisssues such as the lung, liver and adipose tissue. Thus, neutrophilia was detectable in blood and tissues of C26 mice during cancer progression and in pre-cachectic KPC mice compared to PC controls (Fig 1G-J) resulting in an elevated NLR in the circulation (Fig EV1F).

Consistent with the early migration of neutrophils into non-cancerous tissues, we detected increased myeloperoxidase (MPO), a neutrophil marker, in the hepatic and pulmonary parenchyma of pre-cachectic KPC mice, in areas where no microscopic metastatic growth was detectable (Fig EV1G, I). Furthermore, gene expression analysis showed increased levels of neutrophil chemokines in the livers and lungs of KPC mice, including *Cxcl1, Cxcl2* and *Cxcl5* (Fig EV1H, J); corresponding increased expression of *Cxcr2* was observed in both organs, while Cxcr1 was marginally increased in the lung. In line with flow cytometry data, we did not see an increase in the monocyte chemokine *Ccl2.* Liver transaminases were unchanged in pre-cachectic KPC mice (Fig EV1K, L), suggesting that neutrophil infiltration in the liver is not accompanied by signs of hepatocellular damage. No changes in circulating lipids, glucose, or insulin levels were observed (Fig EV1M–P).

Taken together, these findings show the dynamics and spatial distribution of immune changes that precede cachexia onset. Notably, a quantifiable neutrophilia is detectable in established murine models of inflammation-associated epithelial cancer at an early time point during cancer progression, when the organism is otherwise seemingly unaffected by the tumor.

### Pre-cachectic mice exhibit increased BM and splenic extramedullary granulopoiesis

We next performed FACS quantification to investigate the abundance of immune progenitor populations in the spleen and bone marrow, the main hematopoietic organs. Compared to control PC mice, both long-term (LT) and short-term (ST) hematopoietic stem cell (HSC) populations were increased in the spleen of pre-cachectic KPC mice, along with increased multipotent progenitor and granulocyte-monocyte progenitor populations; in contrast, only the LT-HSC population was increased in the bone marrow (Fig 2A, B). In the spleens of pre-cachectic KPC mice, we observed a significant increase in immature neutrophils/late progenitor cells, namely metamyelocytes and band cells (Fig 2C–E). Neutrophil late progenitor cell levels were also increased in the blood and bone marrow, though to a lesser extent (Fig 2F, G). The increased spleen size and abundance of splenic HSC and neutrophil progenitor cells suggest that splenic hematopoiesis is the primary origin of systemic neutrophilia in pre-cachectic KPC mice.

**Fig2.**
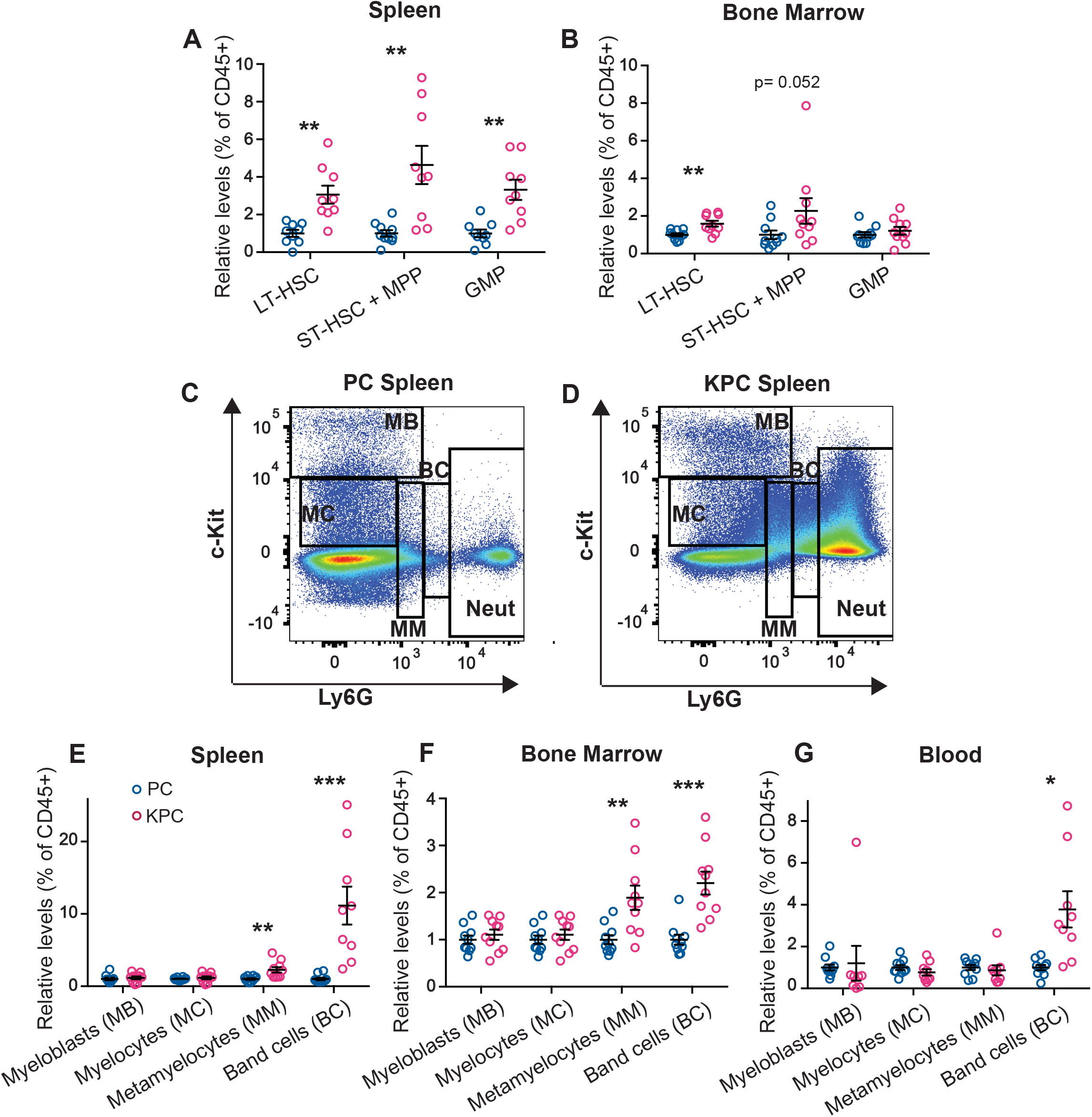
Quantification and FACS gating of progenitor cells, neutrophil precursors, and neutrophil populations in the KPC model. (A-B) Quantification of HSC and progenitor cells (displayed as % out of all CD45+ cells) in the spleen (A) and bone marrow (B) of pre-cachectic KPC and PC controls; (C-D) FACS gating strategy for neutrophils and neutrophil precursor populations in the spleen of PC controls (C) and pre-cachectic KPC mice (D); (E-G) Quantification of neutrophil precursor populations (displayed as % out of all CD45+ cells) in the spleen (E), bone marrow (F) and blood (G) of pre-cachectic KPC mice and PC controls.

### Neutrophils display an inflammatory transcriptional profile and distinctive aerobic glycolytic metabolism during cancer progression

Given that systemic neutrophilia precedes onset of cachexia, we decided to investigate the transciptome of neutrophils in pre-cachectic KPC or PC litermate control mice. Unsupervised differential gene expression analysis demonstrated two distinct clusters comprising those neutrophils isolated from cancer-bearing animals and those from littermates that did not have cancer (Fig 3A). KPC-derived neutrophils exhibited elevated expression of pro-inflammatory genes (Il1b*, mmp8*/9, S100a8/a9), ROS-mediated pathogen killing (Ncf1/2/4, Rac1/2) and genes associated with neutrophil extracellular trap (NET) formation (Ctsb, Ctsd). In terms of maturation-associated genes, KPC-derived neutrophils exhibited elevated Itgam, Tlr4, Cd44, Cd24 relative to PC-derived neutrophils. We next performed Ingenuity pathway analysis of the differentially expressed genes; relative to neutrophils from PC littermate controls, those from pre-cachectic KPC mice exhibited activation of cytokine-driven pathways, cytoskeleton remodeling, and enhanced glycolysis. This is consistent with activation of the upstream regulators IL-6 (a cytokine that is tumor associated in C26 and KPC mice), interferon gamma (IFNγ), IL-17α, tumor necrosis factor (TNF)^18^, IL-1β, and granulocyte colony stimulation factor (G-CSF), as well as Hypoxia-Inducible Factor 1 alpha (HIF-1α), the oxygen-sensitive master regulator of glycolysis under normoxic conditions (Fig 3B). Moreover, *Hif1a* and hallmark hypoxia genes were were significanly overexpressed in KPC-derived neutrophils. Gene set enrichment analysis (GSEA) further identified increased expression in neutrophil degranulation, activation, apoptosis, pentose phosphate pathway (PPP), and glycolysis genes relative to PC-derived neutrophils. Conversely, PC-derived neutrophils were enriched for genes involved in myeloid cell development, fatty acid metabolism and oxidative phosphorylation (Fig EV2A-B). These data suggest that neutrophils from pre-cachectic mice undergo activation-induced metabolic changes.

**Fig3.**
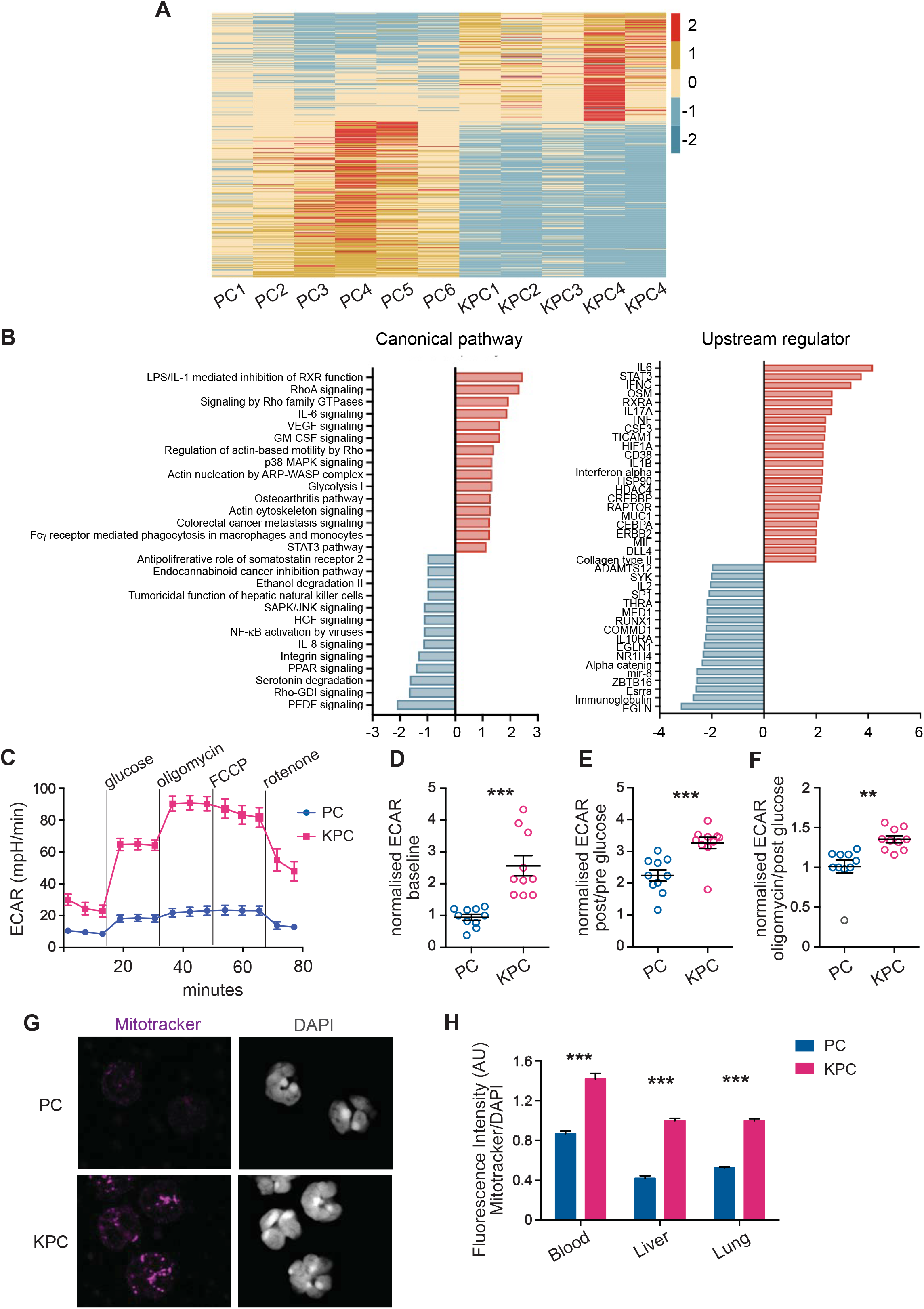
Characterization of immune cell metabolism in pre-cachectic KPC mice. (A-B) Unsupervised differential gene expression analysis (A) and upregulated canonical pathways and upstream regulators (B) from Ingenuity pathway analysis on differentially expressed genes in sorted circulating neutrophils from pre-cachectic KPC mice and PC controls. (C) Extracellular acidification rate (ECAR) of circulating leukocytes from pre-cachectic KPC mice and PC controls measured in real-time by Seahorse assay at baseline, after glucose administration, and after treatments with oligomycin, phenylhydrazone (FCCP), and rotenone; (E-G) Normalized ECAR measurements (ratio of compared timepoints) in circulating leukocytes from mice in Figure 3D at baseline (D), after glucose administration (E) and oligomycin treatment (F); (G) Representative MitoTracker and DAPI immunofluorescence staining in sorted circulating neutrophils from pre-cachectic KPC mice and PC controls; (H) Quantification of the fluorescence intensity (AU) of MitoTracker staining relative to DAPI staining in circulating, hepatic, and pulmonary neutrophils from mice in Figure 3C.

We next analyzed the metabolic phenotype of these activated neutrophils in pre-cachectic mice. Neutrophils are, as first cellular responders of the innate immune system, very easily activated during sample processing, which can result in cell clumping and affect neutrophil quality and quantity^19^. In order to preserve the metabolic profile of the cells, and because the leukocyte fraction was mostly comprised of neutrophils in KPC mice, we used total leukocytes at this point. Glycolytic flux, measured by the extracellular acidification rate (ECAR), was significantly elevated in circulating leukocytes from pre-cachectic KPC mice: at baseline; after glucose administration; and after independent treatments with the oxidative phosphorylation inhibitor oligomycin (a potent disruptor of ATP synthesis), with phenylhydrazone (FCCP), and with rotenone (an inhibitor of the electron transport chain in mitochondria) (Fig 3C-F). Oxygen consumption rate (OCR) was also increased in leukocytes from pre-cachectic KPC mice (Fig EV2C, D). We then examined whether circulating and tissue-infiltrating neutrophils were responsible for the observed changes. Neutrophils from pre-cachectic KPC mice exhibited different mitochondrial activity than those from control PC mice. Quantification of mitochondrial mass using MitoTracker (Deep Red FM) showed increased mitochondrial mass in hepatic, pulmonary, and circulating neutrophils of KPC mice (Fig 3G, H). In line with the enhanced mitochondrial metabolism observed in leukocytes, isolated circulating neutrophils from KPC mice showed increased staining for carnitine palmitoyltransferase 1 (CPT1), the enzyme responsible for transferring fatty acids inside the mitochondria (Fig EV2E, F). Thus, neutrophils from pre-cachectic mice exhibit an increase in overall metabolism and reliance on glycolysis.

The metabolic switch to aerobic glycolysis, termed the Warburg effect, has been observed in both activated immune cells and tumors. Here, we find that this transition characterizes neutrophils during cancer progression. Combined with the known utilization of Warburgian metabolism in cancer cells, these findings indicate that this metabolic state concurrently exists in both cell types within the same organism during cancer progression. Our results suggest that neutrophilia in cancer progression is an energy-expending process and may impact metabolic homeostasis systemically. To understand the relevance of aerobic glycolysis in the context of cachexia, we next targeted this pathway.

### Targeted inhibition of aerobic glycolysis leads to reduced tumor size and expands widespread neutrophilia

Inhibiting aerobic glycolysis to down-modulate the immune response is a therapeutic strategy currently employed for psoriasis and multiple sclerosis^20^. Given the relevance of modified glucose metabolism to activated neutrophils and cancer cells, we assessed whether modulating aerobic glycolysis could affect cancer progression and the clinical severity of cancer-associated cachexia.

Glyceraldehyde 3-phosphate dehydrogenase (GAPDH), the glycolytic enzyme that is rate-limiting exclusively in cells that rely on aerobic glycolysis, can be selectively inhibited by heptelidic acid^20–23^. Pre-cachectic C26 tumor-bearing mice and controls were treated daily with heptelidic acid or vehicle from day 14 post-subcutaneous injection of C26 cells. Treatment with heptelidic acid in C26 pre-cachectic mice resulted in significantly reduced tumor growth rates relative to C26 tumor-bearing mice administered vehicle (Fig 4A), confirming the relevance of aerobic glycolysis to cancer progression in this model. However, overall survival was reduced in the C26 tumor-bearing mice receiving heptelidic acid, while the same treatment did not impact survival of control littermates (Fig 4B). Body composition analysis showed no effect of heptelidic acid in control mice. In contrast, C26 tumor-bearing mice treated with heptelidic acid exhibited significant splenomegaly and skeletal muscle loss without changes in white adipose tissue, compared to their vehicle-treated counterparts (Fig 4C–E). It might be expected that inhibiting glycolytic flux in neutrophils would result in immune cell down-modulation^24^, but we observed a further increase in neutrophilia and splenomegaly in pre-cachectic C26 mice following heptelidic acid administration. A possible explanation is that inhibition of glycolysis is known to impair macrophage efferocytosis, the major mechanism of apoptotic neutrophil clearance in peripheral tissues^26^.

**Fig 4.**
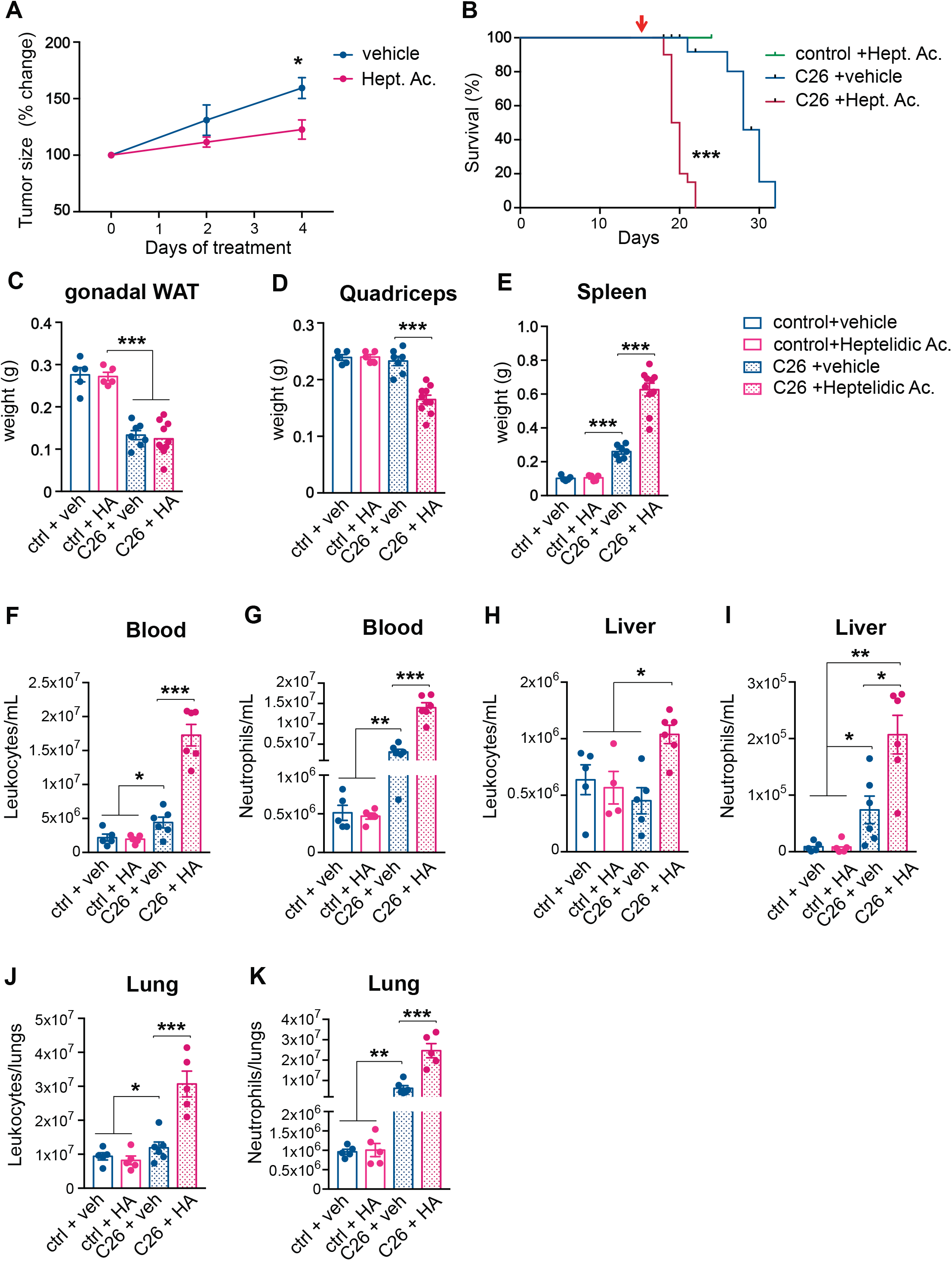
Effects of systemic inhibition of aerobic glycolysis on the severity of cancer cachexia. (A) Longitudinal tumor measurements of C26 mice treated with heptelidic acid or vehicle; (B) Overall survival of C26 mice and littermates treated with heptelidic acid or vehicle; (C-E) Tissue weights of gonadal white adipose tissue (gWAT) (C), quadriceps (D), and spleen (E) of time-matched mice in Fig 4A; (F-K) Quantification of leukocyte (F, H, J) and neutrophil (G, I, K) counts in the circulation (F-G), the liver (H-I), and the lungs (J-K) of mice in Fig 4A.

While immune cell counts of control mice remained unchanged upon treatment, heptelidic acid prompted a further increase in circulating neutrophilia and an increased neutrophilia in lung and liver in pre-cachectic C26 mice, compared to vehicle-treated pre-cachectic C26 mice; leukocyte counts were also elevated in these organs (Fig 4F–K). Our observations thus reveal a therapeutic paradox: systemic inhibition of aerobic glycolysis reduces tumor growth but accelerates cancer-associated cachexia, with a further increase in systemic neutrophilia and decreased survival. Thus, heptelidic acid treatment uncovers a correlation between survival and aerobic glycolysis in immune cells.

### Short-term systemic depletion of neutrophils impairs glucose homeostasis during cancer progression

To ascertain whether neutrophilia represents a generalized response to systemic metabolic stress, we studied immune changes in non-cancer-bearing and cancer-bearing mice subjected to 24-hour total food restriction (TFR). Using this approach to model cachexia, we previously demonstrated that iatrogenic induction of caloric deficiency in the pre-cachectic stage is sufficient to reproduce the functional energetic deficiency that we term ‘metabolic stress’ observed during cachexia in mice and humans^5^.

Non-tumor-bearing mice showed an increase in both circulating and tissue-infiltrating neutrophils after TFR (Fig EV3A–C), consistent with upregulated expression of the neutrophil chemoattractant ligands CXCL2 and CXCL5, overexpression of the activated immune cell surface marker CD36, and downregulated gene expression of the regulator of lipid biosynthesis and adipogenesis sterol regulatory element-binding protein 1 (SREPB1c), and fatty acid synthase (FAS) in the livers of fasted mice (Fig EV3D). These data suggest a generalized, cancer-independent response of neutrophil proliferation, mobilization, and infiltration to conditions of metabolic stress, consistent with previous reports^30^.

To assess the systemic consequences of neutrophil depletion, we injected mice with an anti-Gr1 depleting antibody: a rat IgG2b antibody that recognizes Ly6G and Ly6C surface markers and depletes neutrophils via the complement-mediated membrane attack complex. We used anti-Gr1 rather than the more neutrophil-specific anti-Ly6G antibody because anti-Ly6G has lower efficiency than anti-Gr1 with regard to depleting neutrophils^27–29^.

Anti-Gr1 administration depleted neutrophils efficiently but displayed a short window of efficacy, with a compensatory rebound in neutrophil levels after a few days of treatment, despite repeated administration every 48 hours (Fig EV3E, F). Therefore, studying the effect of long-term pharmacological neutrophil depletion on cancer progression and cachexia was not experimentally possible using this model. Anorexia represents a key pathologic event in clinical cachexia, thus we used this short-term depletion window to assess the contribution of neutrophils to the metabolic consequences of TFR.

Non-tumor-bearing mice as well as pre-cachectic C26 tumor-bearing mice were treated with anti-Gr1 antibody or an isotype control and were challenged with TFR or allowed free access to food (Fig EV3G). Body weight loss after TFR was similar in the control and pre-cachectic C26 mice, and it was not affected by anti-Gr1 treatment (Fig EV3H). However, body weight recovery post-TFR was impaired in anti-Gr1-treated C26 mice compared to isotype-treated C26 mice and controls, which fully recovered their original weight after 24 hours of refeeding, suggesting dysfunctional macronutrient handling in the neutrophil-depleted, fasted C26 mice (Fig 5A, B). Although serum glucose levels were similar in non-tumor bearing controls and pre-cachectic C26 mice in the fed state or after TFR (Fig 5C), neutrophil depletion caused hypoglycemia in pre-cachectic C26 mice subjected to TFR (Fig 5D).

**Fig 5.**
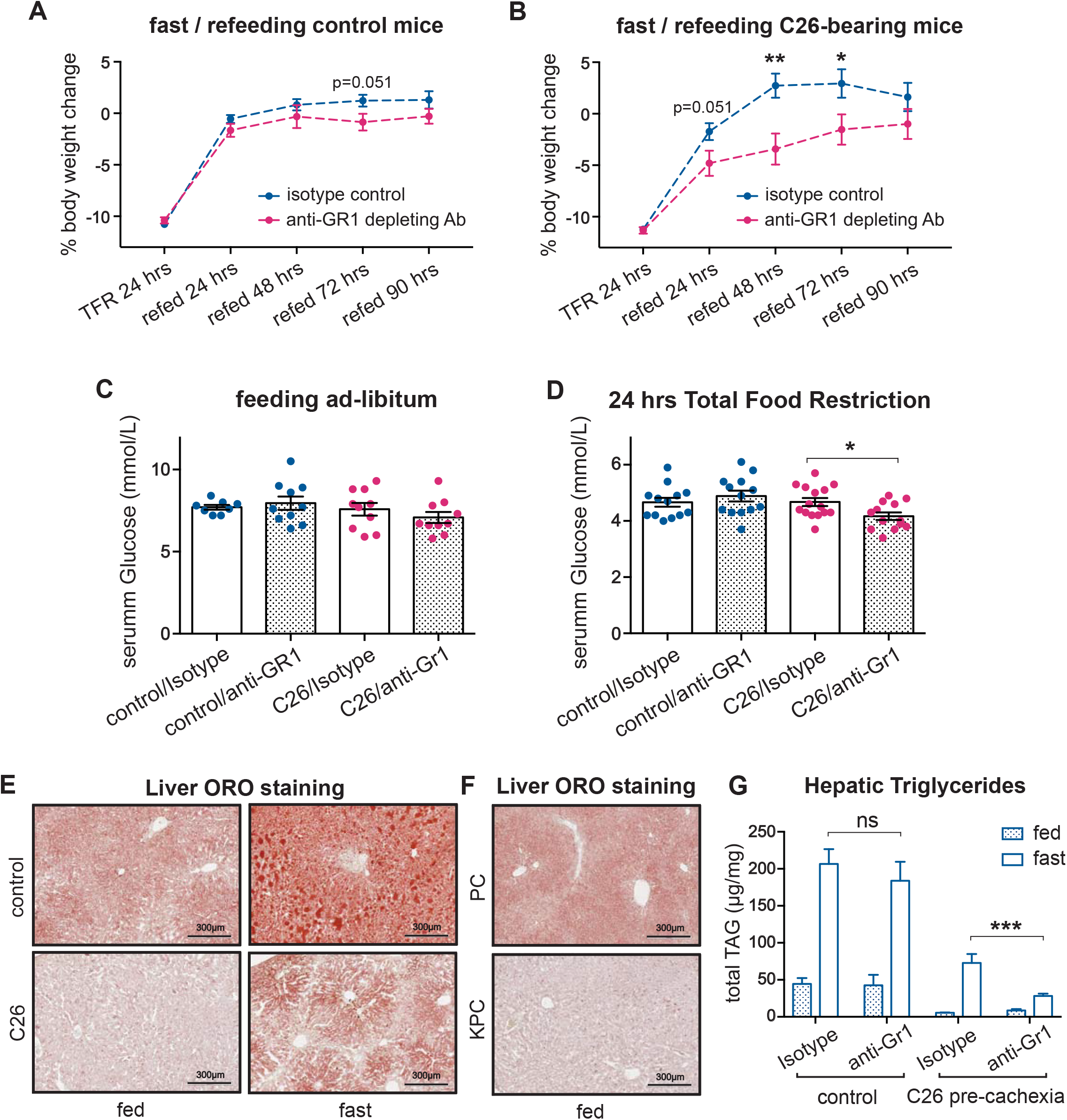
Effect of acute neutrophil depletion during cancer-associated metabolic stress. (A-B) Body weight trajectories after TFR of littermates (A) and pre-cachectic C26 mice (B) treated with anti-Gr1 or isotype. (C-D) Glucose levels in C26 mice and littermates treated with anti-Gr1 or isotype, when fed *ad libitum* (C) and after 24 hours of total food restriction (TFR) (D); (E-F) Lipid staining by Oil Red O in the liver of fed or fasted pre-cachectic C26 mice and littermates (E) and fed pre-cachectic KPC mice and PC controls (F); (G) Quantification of triglycerides by liquid chromatography-mass spectrometry in the liver of fed and fasted pre-cachectic C26 mice and littermates treated with anti-Gr1 or isotype;

WAT lipolysis is essential for providing lipid substrates for hepatic gluconeogenesis, which sustains normal serum glucose levels during fasting^31^. Therefore, the hypoglycemic phenotype observed in the neutrophil-depleted pre-cachectic mice led us to investigate hepatic lipid metabolism. Lipid staining showed the expected physiological accumulation of hepatic lipids upon TFR in both controls and pre-cachectic C26 mice; however, the intensity was visibly reduced in pre-cachectic C26 mice, both in the fed and TFR states (Fig 5E). Similarly, a decrease in hepatic lipid accumulation was observed in the livers from pre-cachectic KPC mice compared to control PC mice (Fig 5F). No obvious difference in hepatic glycogen staining was observed (Fig EV3I). Quantification of lipid triglycerides confirmed that levels were reduced in livers from pre-cachectic C26 mice, both in the TFR and the fed states. Remarkably, while neutrophil depletion did not affect hepatic triglyceride levels in nontumor bearing controls, it significantly reduced triglyceride levels in fasted pre-cachectic C26 mice (Fig 5G). Of relevance, neutrophils from pre-cachectic mice display increased mRNA expression levels of lipolytic cytokines, e.g., TNFα and IL-1β^32,33^ and enzymes that modify glycemic indexes, such as GPLD1^34^ (Fig EV3J). Collectively, these data support a model wherein neutrophil depletion reduces the availability of hepatic lipids, thereby impairing gluconeogenesis in cancer-bearing mice during conditions of acute metabolic stress, such as TFR. Thus, our observations indicate that cancer-associated neutrophilia and inflammation-induced neutrophil activation modulate substrate supply for systemic glucose homeostasis in cancer progression. These findings further support the notion that neutrophilia acts as an adaptive/protective mechanism that maintains systemic metabolic homeostasis during cancer progression.

## DISCUSSION

Cancer is a progressive disease that is initiated by cellular changes that promote tumor formation and often ends with death of the organism. Overt cachexia represents an irreversible late stage of the disease and results from a sequence of metabolic events in cancer progression, including lean body mass wasting, hypercatabolism, weight loss and insulin resistance. Since cancer-associated cachexia is a clear manifestation of the ongoing interactions between the tumor and the host metabolism, these interactions must occur throughout the stepwise process of cancer progression.

We, thus, identify neutrophilia and aerobic glycolysis by neutrophils as an early display of tumor-host reciprocity that when targeted, both cellularly or metabolically, worsens outcome and accelerates cancer progression. This demonstrates an actively adaptive or protective response of the host against the advancing tumor that aims to sustain its metabolic homeostasis. Persisting efforts to achieve physiological balance may later on become depleting and therefore trigger irreversible consequences in cancer progression, like cancer-associated cachexia.

Regarding the translation of these findings, we note that neutrophilia in our *in vivo* models was first detectable in solid organs such as liver and lung, most likely due to active neutrophil recruitment, and only later in the circulating blood, which is the preferred medium of biomarker detection in the clinic. The later appearance of neutrophilia in the circulation may therefore falsely suggest a later onset of systemic dysregulation. In agreement with recent literature^35^, we found that cancer-associated neutrophilia is mostly derived from increased splenic hematopoiesis, through expansion of HSC and progenitors. Mechanistically, it is driven by local recruitment and accumulation of rare extramedullary HSC and progenitor cells in the splenic red pulp, mediated by the CCL2/CCR2 axis^36^. The spleen is known to contribute significantly to hematopoiesis in a range of inflammatory states^37–39^, suggesting that this is an adaptive response to chronic inflammation. The factors driving neutrophila in early disease remain unclear, although recent work suggests that primary tumour-driven ‘leaky gut syndrome’ can contribute to systemic inflammation. Expression of neutrophil chemokines in peripheral organs, and the activated transcriptomic signatures of pre-cachectic mouse neutrophils further support the hypothesis that primary tumours trigger microbial-driven activation.

Such a putative protective role for neutrophils in cancer progression has diverse clinical implications, because therapeutic agents given to patients with cancer frequently alter neutrophil counts. Chemotherapy, for example, can cause life-threatening neutropenia as a side effect, and blocking neutrophil migration via CXCR1 and CXCR2^40^, and inhibiting G-CSF^41^, inhibit neutrophil function. Our model helps to partly explain the high mortality from neutropenic sepsis in patients with cancer^42^, as infection and neutropenia would synergize to promote metabolic stress. On the other hand, steroids, G-CSF, and granulocyte-macrophage colony-stimulating factor (GM-CSF) increase neutrophil counts and our study suggests that this may have effects on systemic metabolic homeostasis.

A recent study showed that infiltration of neutrophils in the brain worsens cachexia severity^43^ and future work should aim to delineate the contribution of organ-specific neutrophil infiltration. Expression of genes involved in NET formation and activation status in neutrophils from KPC mice is in agreement with recent literature^44,45^. In this work, we focused on the definition of the systemic role of neutrophils in the pathophysiology of cachexia. While the intrinsic cellular consequences of metabolic modulation have been thoroughly explored in neutrophils^8^, their impact on host metabolism was unknown until now. We find that neutrophils contribute to the hosts’ metabolic homeostasis during cancer progression and, thereby, suggest that treatments affecting neutrophils will have complex and potentially detremental effects on the host. These results argue for the absolute necessity of a combined analysis regarding the effects of candidate cancer treatments on the tumor and the host.

Treatment effectiveness is often assessed using surrogate outcomes, such as reduction in tumor size or increased time until radiological progression, termed “progression-free survival”^46^, but it is increasingly recognized that these surrogate measures are poor predictors of overall survival^47^. Our results demonstrate that the host response is an important determinant of cancer outcome and may thus serve as an explanation of the insufficiency of tumor assessment to predict outcome. In this context, our finding that GAPDH inhibition leads to tumor regression but accelerated death is particularly striking. As heptelidic acid should act only on tumor cells and immune cells, in particular activated neutrophils^20^, this divergent phenotype indicates that the effect of GAPDH down-modulation on immune cells in the host outweighs its inhibitory effect on tumor growth. Targeting of upstream molecules, like GLUT1, show similar effects in inhibiting tumor growth but may cause normal cell death because they prevent oxygen transport and endothelial cell angiogenesis for red blood cells^48,49^. These findings provide a powerful proof-of-concept that paradoxical tumor-host responses to treatment occur *in vivo*, warranting further investigation in the context of other treatments^50^.

Our findings highlight the importance of distinguishing adaptive from detrimental processes during cancer progression. The assumption that reversal of a process observed during cancer progression will result in improved outcome may not hold true. In fact, we demonstrate that targeting metabolic pathways that are relevant to both host and cancer cells may lead to worsening of outcome despite tumor regression.

## MATERIALS AND METHODS

### Laboratory animals

Two different mouse models of cachexia were used: a transplanted C26 model of colorectal cancer and the genetically engineered autochthonous KPC model of pancreatic cancer. The C26 model is based on wild-type BALB/c mice that are inoculated subcutaneously with a syngeneic tumor. In the KPC system, an activating point mutation (G12D) in Kras and a dominant negative mutation in Trp53 (R172H) are conditionally activated in the pancreas by means of Cre-Lox technology. Both preclinical models have been shown to develop tumors that secrete IL-6, therefore the host is unable to produce ketones during the caloric deficiency associated with cachexia, causing a rise in glucocorticoid levels. KPC and BALB/c mice were obtained from Charles River Laboratories. All mice were kept in pathogen-free conditions on a 24-hour 12:12 light-dark cycle and allowed to acclimatize for 7 days. For pharmacological inhibition of aerobic glycolysis, C26 syngeneic colon cancer mice and control mice received daily intraperitoneal injections with heptelidic acid (1 mg/kg; Cayman Chemical). For anti-IL-6 treatment, C26 tumor-bearing mice received intraperitoneal injections with InVivoMAb anti-mouse IL-6 (1 mg in 200μl PBS; Bio X Cell, clone MP5-20F3). Body weight, food intake, and clinical signs were monitored on a daily basis. Handling was kept to a minimum. Mice were defined as cachectic and were sacrificed when showing clinical signs of sickness or when their weight loss exceeded 10% peak weight (usually corresponding to the starting body weight at day 0). Death was confirmed by cervical dislocation. Experiments and care of C26 syngeneic colon cancer mice and control mice, and KPC and PC were performed in accordance with national and institutional guidelines and approved by the UK Home Office, the animal ethics committee of the University of Cambridge.

### Survival studies

Weight-stable, PDAC-bearing KPC mice with tumors 3–5 mm in size and no evidence of obstructive common bowel duct were enrolled in the experiments together with their weight- and age-matched PC controls. Weight-stable wild-type male BALB/c mice were inoculated subcutaneously in their right flank with 2×10^6^ viable C26 colorectal cancer cells in RPMI vehicle at 100 μl per mouse. C26-injected BALB/c mice were enrolled in the study together with their respective weight- and age-matched non-tumor-bearing BALB/c control littermates.

### Neutrophil depletion

C26 tumor-bearing mice were injected intraperitoneally with 200 μg of GR1-depleting antibody (Ultra-LEAF anti-mouse-Ly6G/Ly6C Ab, BioLegend). Control mice were injected intraperitoneally with 200 μg of isotype control (Ultra-LEAF rat IgG2b, BioLegend). Mice were injected once with GR1-depleting antibody or isotype control on day 12 post C26 cancer cell subcutaneous injection.

### Total food restriction (TFR)

Mice were singly-housed and moved into new cages containing only bedding, nesting, material and a water bottle. TFR was initiated in the middle of the 12-hour light period at 12:00pm, and mice were re-fed after 24 hours.

### Tumor size

PDAC tumors were detected via palpation and confirmed by high-resolution ultrasound imaging (Vevo 2100, VisualSonics), and ultimately at necropsy. Tumor growth was monitored by ultrasound scans assessed at multiple angles. Mice were carefully observed for any macroscopic metastases. Tumor development in BALB/c mice was determined via palpation and monitored daily by caliper measurements. The maximum crosssectional area and maximum diameter of the tumors were determined for each time point.

### Plasma and serum analysis

Tail bleeds and terminal cardiac bleeds were taken. Tail vein bleeds were performed using a scalpel via tail venesection without restraint, and terminal bleeds were obtained through exsanguination via cardiac puncture under isoflurane anesthesia. Samples were kept on ice at all times. Plasma samples were collected into heparin-coated capillary tubes to avoid coagulation and were processed as follows: centrifuge spin at 14,000 rpm for 5 minutes at 4°C, snap frozen in liquid nitrogen, and stored at −80°C. Analyses of alanine aminotransferase (ALT), aspartate aminotransferase (AST), triglycerides, free fatty acids (FFA), glucose, and insulin were performed by the Core Biochemical Assay Laboratory, University of Cambridge.

### Tissue collection

PDAC tumors, liver, spleen, quadricep muscle, subcutaneous fat, and lungs were collected and weighed during necropsy dissection. PDAC tumors, liver, spleen, quadricep muscle, subcutaneous fat, lungs, and hypothalamus were collected and weighed during necropsy dissection. Subsequently, tumor, liver, and spleen samples were cut into two equal parts, which were either snap frozen in liquid nitrogen or fixed in 10% neutral buffered formaldehyde for 24 hours at room temperature before being transferred to 70% ethanol and later paraffin-embedded or immunohistochemistry processing. All the other organs and tissue samples were immediately snap frozen and stored at −80°C.

### RNA isolation, reverse transcription, and quantitative RT-PCR (qRT-PCR)

Frozen tissue samples were lysed in 1 ml of TRIzol (Sigma) using a Precellys^®^ tissue homogenizer. A total of 200 μl of chloroform was added, and the samples were centrifuged for 15 minutes at 13,200 rpm at 4°C. The aqueous phases containing the RNA were carefully transferred to new tubes, and 500 μl of isopropanol were added for precipitation of the RNA. After vigorously shaking, the samples were centrifuged for 30 minutes at 13,200 rpm at 4°C. Supernatants were removed, and the samples were washed by adding 1000 μl of cold 80% ethanol followed by centrifugation at 13,200 rpm for 10 minutes at 4°C. Once dried, the pellets were dissolved in nuclease-free water; the added volume depended on the pellet size. Then the samples were stored at −80°C until they were used. To purify RNA samples, DNAse treatment was performed: 1–2 μg of RNA were diluted in a total volume of 10 μl, including 1 μl of reaction buffer and 1 μl of DNAse I (1U/ μl), and nuclease-free water. After incubating for 15 minutes at room temperature, 1 μl of 25 mM EDTA was added to inactivate the reaction. A total of 10 μl of GoScript™ Reverse Transcription Mix was prepared for each cDNA reaction (4 μl of nuclease-free water, 4 μl of GoScript™ Reaction Buffer, Oligo (dT), and 2 μl of GoScript™ Enzyme Mix, Promega). This mastermix was combined with the RNA sample (final volume of 20 μl) and, after mixing well, the reaction was incubated following these steps: 1) anneal primer (5 minutes at 25°C), 2) extension (60 minutes at 42°C), and 3) inactivation (15 minutes at 70°C). Samples were stored at −20°C until qPCR was performed. The Quantitative PCR Primer Database (QPPD) was used to search for primers previously validated in the literature (https://pga.mgh.harvard.edu/primerbank/). The NCBI Standard Nucleotide BLAST online tool was used to verify the primers (https://blast.ncbi.nlm.nih.gov/Blast.cgi?PAGE_TYPE=BlastSearch). A total of 2 μl of cDNA, 5 μl of Sybr Green qPCR Master mix, 3 μl of nuclease-free water, and 0.2 μl of primers (0.1 μM) were used per reaction. The comparative cycle threshold method was used for quantification, and expression levels were normalized using the housekeeping gene β-actin. All samples were run in technical triplicate. The primer sequences are available upon request.

### Single cell preparation

Cell suspensions were prepared from tissues by mechanical dissociation, followed by digest in 5 ml of RPMI-1640 containing collagenase I (500 U/ml) and DNase I (0.2 mg/ml) for 45 minutes at 37°C on a shaker (220 rpm), followed by filtration through a 70-μm strainer and 25% Percoll gradient enrichment of leukocytes, and red blood cell (RBC) lysis. Tumor cells were recovered without Percoll enrichment. Blood cells were lysed in 5 ml of RBC lysis buffer three times for 5 minutes, and spleens were strained through a 70-μm filter in RPMI-1640 before lysing erythrocytes with RBC lysis buffer for 5 minutes. Single cells were restimulated and stained for surface and intracellular markers (see flow cytometry).

### Flow cytometry

Single cells were incubated with anti-mouse CD16/32 (Thermo Fisher) to block Fc receptors and stained as indicated. The lineage cocktail contained ±CD3 (145-2C11), ±NK1.1 (PK136), TCRβ, CD5 (53-7.3), CD19 (1D3), CD11b (M1/70), CD11c (N418), FcεR1α (MAR-1), F4/80 (BM8), Ly-6C/G (Rb6-8C5), and Ter119 (TER-119) all on eFluor450 (eBioscience). For intracellular staining, we used the Foxp3/Transcription Factor Kit (Thermo Fisher) or Cytofix/Cytoperm Kit (BD Biosciences) as per the manufacturers’ instructions. For intracellular cytokine detection, single cells were stimulated with PMA (60 ng/ml) and ionomycin (500 ng/ml) plus 1× protein transport inhibitor (Thermo Fisher), 1× cytokine stimulation cocktail (Thermo Fisher), or platebound anti-NK1.1 mAb (10–30 μg/ml, BioXcell), or recombinant IL-12 (20 ng/ml) and IL-18 (5 ng/ml) in culture media (RPMI-1640, 10% FCS) at 37°C for 3 hours before staining. Data were acquired on a BD Fortessa or Symphony instrument (BD Biosciences), and cells were quantified using CountBright beads. Data were analyzed using FlowJo X (Tree Star). B220 (RA3-6B2, Life Technologies, APC.eFl780), CD3e (145-2C11, eBioscience, PE.Cy7 and eFl450) (25-0031-83, 4304567), CD4 (RM4-5, eBioscience, AF700), CD5 (53-7.3, eBioscience, eFl450), CD8 (53-6.7, eBioscience, PerCP.eFl710 and SB645), CD11b (M1/70, eBioscience, eFl450, APC.eFl780, and BV785), CD11c (N418, eBioscience, eFl450 and AF700), CD16/32 (93, BioLegend), CD19 (1D3, eBioscience, eFl450), CD31 (390, BioLegend, BV605), CD45 (30-F11, BioLegend, BV510), CD64 (X54-5/7.1 BioLegend, BV711), CD127 (SB/199, BD Biosciences, PE.CF594), CD172a (P84, BioLegend, AF488), c-Kit (BioLegend), EpCam (G8.8, BioLegend, BV711), FceR1a (MAR-1, eBioscience, eFl450 and PerCP.eFl710), Fixable Viability Dye (eBioscience, UV455), F4/80 (BM8, eBioscience, eFl450 and APC.eFl780), I-A/I-E (CI2G9, BD Biosciences, BUV395), Ly-6C/G (Rb6-8C5, eBioscience, eFl450), Ly-6G (1A8-Ly6g, eBioscience, PE.eFl610), Ly-6C (HK1.4, eBioscience, PE.Cy7), NK1.1 (PK136, BD Biosciences, BUV395 and eBioscience, eFl450), RELMα (DS8RELM, Invitrogen, APC), Podoplanin (8.1.1, BioLegend, PE.Cy7), Roryt (Q31.378, eBioscience, PerCP.eFl710), SiglecF (1RNM44N, eBioscience, SB600), Ter119 (TER-119, eBioscience, eFl450).

### Histology

Lung and liver lobes were fixed in 10% neutral-buffered formalin in phosphate-buffered saline (PBS) for 24 hours, followed by transfer to 70% Ethanol in PBS for another 24 hours and embedded into paraffin. Then, 3-μm sections were cut and stained with antimyeloperoxidase antibody (Abcam, ab9535). The CRUK-CI Histology Core performed tissue embedding, sectioning, and staining. Image quantification was performed using the HALO software (HALO, Indica labs). Oil Red O and Periodic Acid-Schiff’s (PAS) stainings were performed by the Metabolic Research Laboratories, University of Cambridge. For Oil Red O staining, fresh frozen sections were fixed in 10% neutral-buffered formalin for 10 minutes, then stained in Stock Oil Red O plus Dextrin solution for 20 minutes and counterstained with Mayer hematoxylin for 30 seconds. For PAS staining, 2–5 μm sections were incubated with Periodic Acid Solution for 5 minutes, then Schiff’s reagent for 15 minutes, and counterstained with hematoxylin for 15 seconds.

### Lipidomics experiments

Liver tissue was homogenized in chloroform/methanol (2:1, 1 mL) using TissueLyser (Qiagen Ltd., Manchester, UK). Deionized water (400 uL) was added and after mixing, a centrifugation step (13,000 x g) was used to separate the organic lipid-containing layer. This was analyzed by liquid chromatography mass spectrometry (LC-MS) on an Accela Autosampler coupled to an LTQ Orbitrap Elite™ (Thermo Fisher Scientific, Hemel Hempstead, UK). Separation was achieved using an Acquity C18 BEH column at 55**°**C. Mobile phase A was 60:40 acetonitrile:water, and mobile phase B was 90:10 isopropanol:acetonitrile, each with 10 mM ammonium formate. A gradient flow at 0.5 mL/min was used, starting with 40% B, to 99% B over 8 minutes, then held at 99% B for 0.5 minutes, then returned to initial conditions. The HESI source conditions were 375**°**C and 380**°**C for source and capillary temperature, respectively; gas flow was 40 arbitrary units. Analysis was performed in positive ion mode by full scan (range *m/z* 200–2000). Triacylglycerol (TAG) identification was performed by accurate mass and retention time using an in-house database. Peak areas were normalized to an isotopically labelled internal standard and tissue weight and summed for total TAG quantification.

### MitoTracker analysis and immunofluorescence

Neutrophils were isolated with a Neutrophil Isolation Kit (Miltenyi Biotec) according to the manufacturer’s instructions. MitoTracker analysis was performed according to the manufacturer’s instructions (Invitrogen, Deep Red FM molecular probes). For immunofluorescence, Anti-CPT1 antibody [8F6AE9] (Alexa Fluor^®^ 488, Abcam, ab171449) was used. Immunofluorescence images were analyzed using Fiji software. Images were binarized, and a mask was created to segment cells in 2D. The segmented cells were saved as regions of interest and were used to measure the mean intensity value of MitoTracker and CTP1. For each neutrophil, a unique averaged fluorescence intensity value was generated.

### RNA sequencing

Libraries were prepared using the NEBNext^®^ Single Cell/Low Input RNA Library Prep Kit for Illumina^®^ (New England Biolabs). Quality control and quantification of the libraries were performed using the Bioanalyzer DNA 1000 kit (Agilent) and the KAPA library quantification kit for Illumina (KAPA Biosystems), respectively. The libraries were then pooled and sequenced on two lanes of the HiSeq 4000 system, Illumina (single end 50).

### Bioinformatics functional analyses

After statistical analysis, differentially expressed genes within groups were studied using Ingenuity Pathway Analysis (IPA, Qiagen). We uploaded in IPA the whole transcriptome and, to run the analyses, the data were filtered for expression abundance (>10 Basemean), statistical significance [padj<0.001(WBC); padj<0.05(NF)], and fold change −1 < Log2FC > 1. “Canonical Pathways” and “Upstream Regulators” networks, describing interactions and relationships experimentally confirmed between differentially expressed genes and others that functionally interact with them, were generated and ranked in terms of the significance of participating genes (p<0.05) and activation status (Z-score).

### Oxygen consumption and extracellular acidification rates measurement

To assess the oxygen consumption rate (OCR) and extracellular acidification rate (ECAR), each well of a XFe24 (Agilent) Cell Culture microplate was coated with 25 μL of a solution containing 22.4 μg/mL CellTak (Corning). The solution was allowed to react for 20 minutes at room temperature and was either used immediately for experiments or stored at 4°C. A coated microplate was pre-warmed at 37°C and 3×10^5^ cells were seeded in 675 μL of bicarbonate-free Dulbecco’s Modified Eagle’s Medium (DMEM) (Sigma-Aldrich, D5030) supplemented with 1 mM pyruvate, 4 mM glutamine, 40 mM phenol red, and 1% v/v Fetal Bovine Serum (FBS). To eliminate carbonic acid residues from the medium, cells were incubated for at least 30 minutes at 37°C with atmospheric CO_2_ in a non-humidified incubator. OCR and ECAR were assayed in a Seahorse XF-24 extracellular flux analyzer by the addition via ports A–D of 25 mM Glucose (port A), 1 mM oligomycin (port B), 1 mM carbonyl cyanide-p-trifluoromethoxyphenylhydrazone (FCCP, port C), 1 mM rotenone, and 1 mM antimycin A (port D) in the same cell culture medium used for cell seeding. Three measurement cycles of 2-minute mix, 2-minute wait, and 4-minute measure were carried out at basal condition and after each injection.

### Statistical analysis

Data are expressed as the mean ± SEM unless otherwise stated and statistical significance was analyzed using GraphPad Prism 7.03 software. For survival analysis, data were shown as Kaplan-Meier curves, and the log-rank (Mantel-Cox) test was used to assess survival differences. For categorical outcomes comparison (clinical signs at necropsy, metastasis, sex, tumor location), the Chi-square test was used. For quantitative data (organ weights), an unpaired two-tailed Student’s t-test was applied with Welch’s correction (does not assume equal standard deviations; SDs). When comparing more than 2 groups at the same time, one-way ANOVA with Tukey’s correction for post-hoc testing was used. For statistical comparison of quantitative data at different times (body weight), unpaired two-tailed Student’s t-tests were performed at each time point with the Holm-Šidák method correction for multiple comparisons.

## ACKNOWLEDGMENTS

Michele Petruzzelli was supported by the NIHR Clinical Lecturer Fellowship and starter grant for Clinical Lecturers from the Academy of Medical Science, UK. Miriam Ferrer is supported by the “la Caixa” Foundation (ID 100010434) in the framework of the “La Caixa” Fellowship Program under agreement LCF/BQ/AA18/11680037, and from the MRC Cancer Unit with an MRC CU Research Studentship. Tobias Janowitz acknowledges funding from Cancer Research UK (C42738/A24868), The Mark Foundation for Cancer Research (33300111), Cold Spring Harbor Laboratory (CSHL), and developmental funds from CSHL Cancer Center Support Grant 5P30CA045508. Timotheus Halim was supported by a fellowship from The Royal Society and Wellcome Trust (204622/Z/16/Z), and Cancer Research UK (CRUK) funding (A24995). The Wagner laboratory is supported by ERC (ERC**-**AdG 2016 CSI**-**Fun, grant number 741888) and the Medical University of Vienna (MUV). This work was supported by Medical Research Council (MRC) Programme grants MC_UU_12022/1 and MC_UU_12022/8 to Ashok Venkitaraman.

## AUTHOR CONTRIBUTIONS

Conceptualization and Methodology, M.P., M.F., T.J., T.H., and A.R.V.; Contribution to Data Acquisition and Analysis, all authors; Investigation, M.P., and M.F.; Resources, T.J., T.H., E. F. W., and A.R.V; Writing – Original Draft, M.P., M.F., and T.J.; Writing – Review & Editing, all authors.; Supervision, T.J., T.H., and A.R.V.; Project Administration, M.P.; Funding Acquisition, T.J., T.H, and A.R.V.

## THE PAPER EXPLAINED

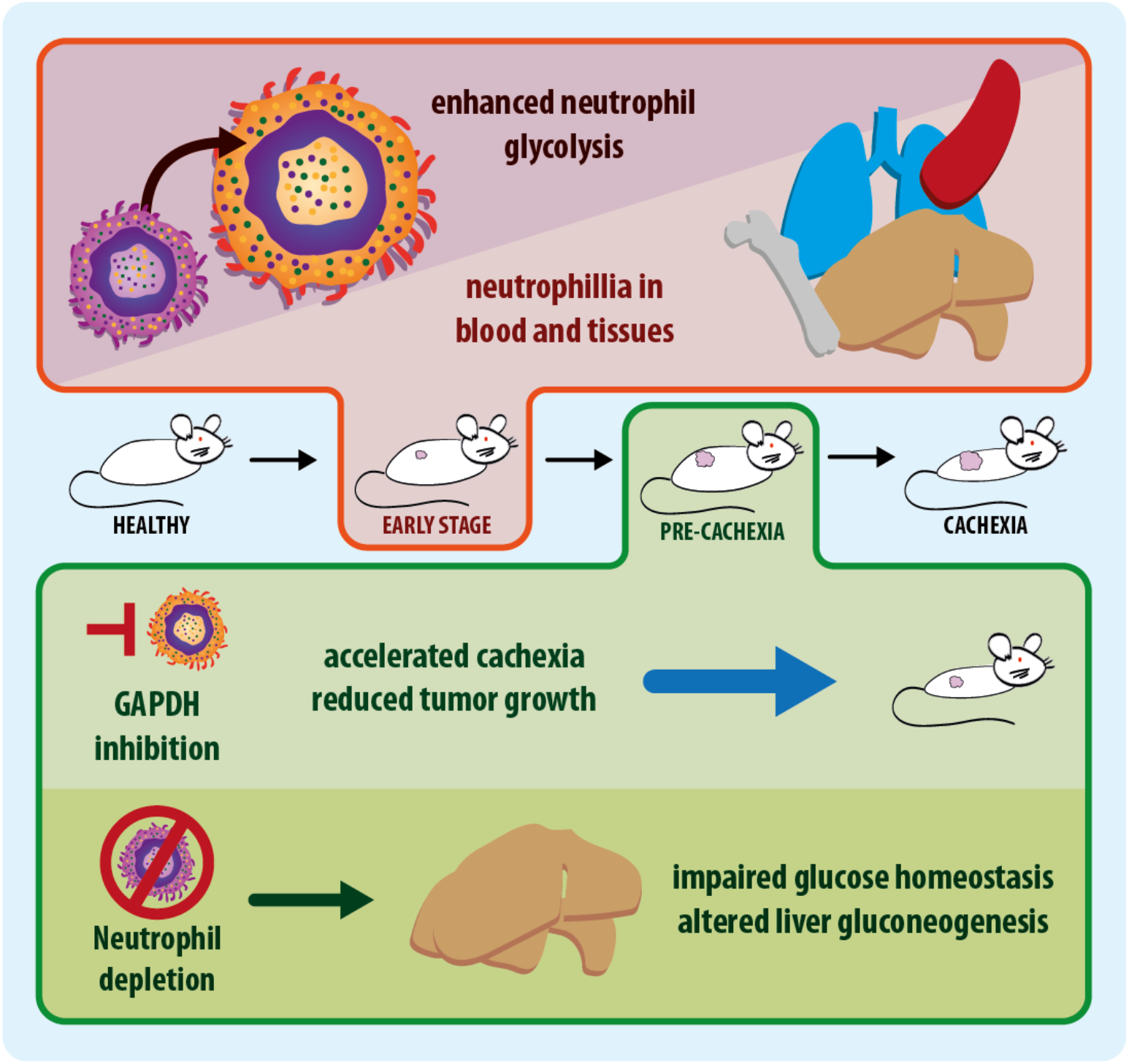

### PROBLEM

Patients with cancer suffer from systemic metabolic impairment during cancer progression, such as hypercatabolism, muscle wasting and insulin resistance. An elevated neutrophil-to-lymphocyte ratio is an early negative predictor of outcome in patients with cancer, but a potential role of neutrophils on the host metabolism has not yet been investigated.

### RESULTS

We identify widespread neutrophilia as an early event in cancer progression, and we find that neutrophils display an enhanced aerobic glycolytic profile. Pharmacological inhibition of aerobic glycolysis, a pathway that also characterizes cancer cells, leads to expanded neutrophilia, reduced tumor size and shorter survival. Quantitative depletion of neutrophils impairs glucose homeostasis and availability of hepatic lipids during cancer progression, suggesting that neutrophils play an adaptive role in metabolic host homeostasis during cancer progression.

### IMPACT

Targeting aerobic glycolysis in cancer uncouples tumor growth from survival through its effect on both tumor and immune cell metabolism. Our results demonstrate that assessment of candidate cancer treatment efficacy should include both tumor and host reponses.

## EXPANDED VIEW FIGURE LEGENDS

**Fig EV1.**
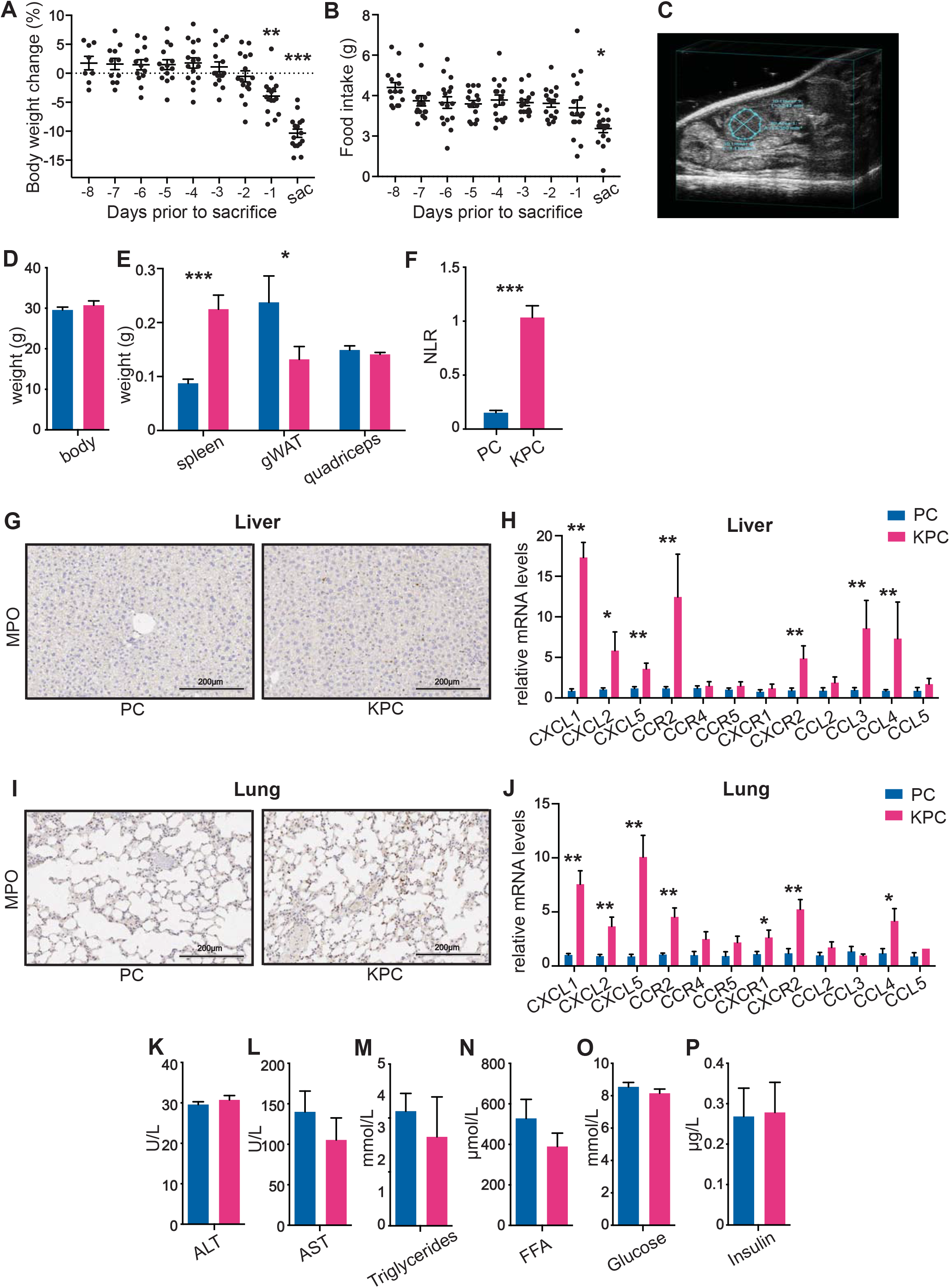
Extended metabolic and immune phenotyping of C26 and KPC models of cancer cachexia. (A-B) Longitudinal change in % body weight (A) and food intake changes (B) during the last 8 days prior to sacrifice in mice injected subcutaneously with C26 colorectal cancer cells; (C) Representative ultrasound image of a pancreatic tumor in a pre-cachectic KPC mouse; (D-E) Body (D), spleen, gonadal white adipose tissue (gWAT) and quadricep (E) weights of pre-cachectic KPC mice and PC littermates; (F) Neutrophil to lymphocyte ratio (NLR) in pre-cachectic KPC mice and PC controls; (G, I) Myeloperoxidase (MPO) staining as marker for neutrophil infiltration in the liver (G) and lung (I) of pre-cachectic KPC mice and PC controls, and relative expression levels of chemokines and chemokine receptors measured by RT-qPCR in the liver (H) and lung (J) of pre-cachectic KPC and control PC mice; (K-P) Quantification of liver transaminases (K-L), triglycerides (M), free fatty acids (FFA) (N), glucose (O), and insulin (P) in the plasma of pre-cachectic KPC mice and PC controls.

**Fig EV2.**
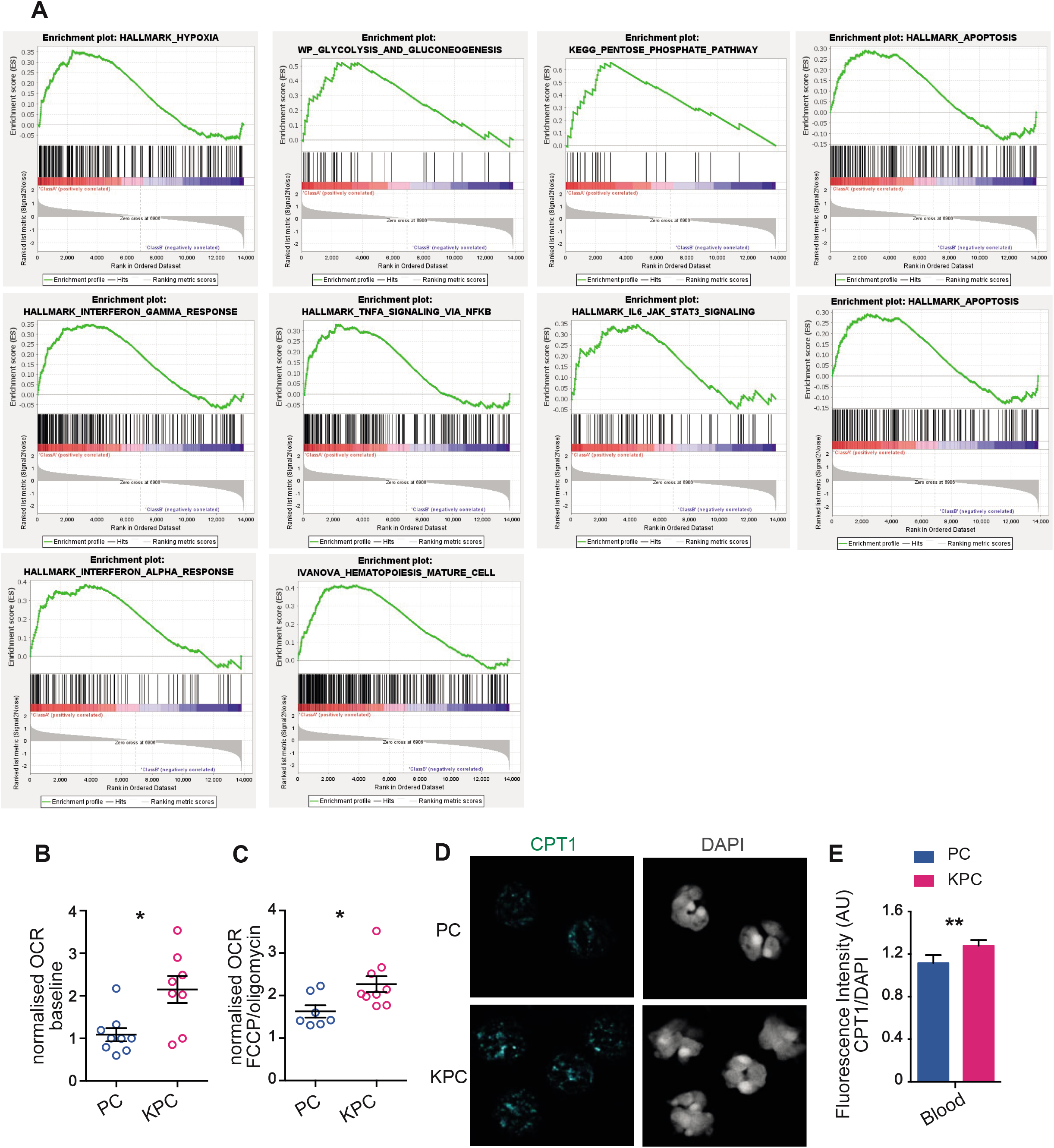
Extended characterization of the transcriptional profile of KPC-derived and PC-derived neutrophils. (A-B) Upregulated genes by Gene set enrichment analysis (GSEA) in KPC-derived (A) and PC-derived (B) neutrophils; (C-D) Normalized oxygen consumption rate (OCR) measurements by Seahorse assay at baseline (D) and after treatment with phenylhydrazone (FCCP) and oligomycin (D) in circulating leukocytes from pre-cachectic KPC mice and PC controls; (E) Representative immunofluorescence staining for carnitine palmitoyltransferase 1 (CPT1) and DAPI in sorted circulating neutrophils from mice in Fig 3A; (F) Quantification of the fluorescence intensity (AU) of CPT1 staining relative to DAPI in sorted circulating neutrophils from mice in Fig 3A.

**Fig EV3.**
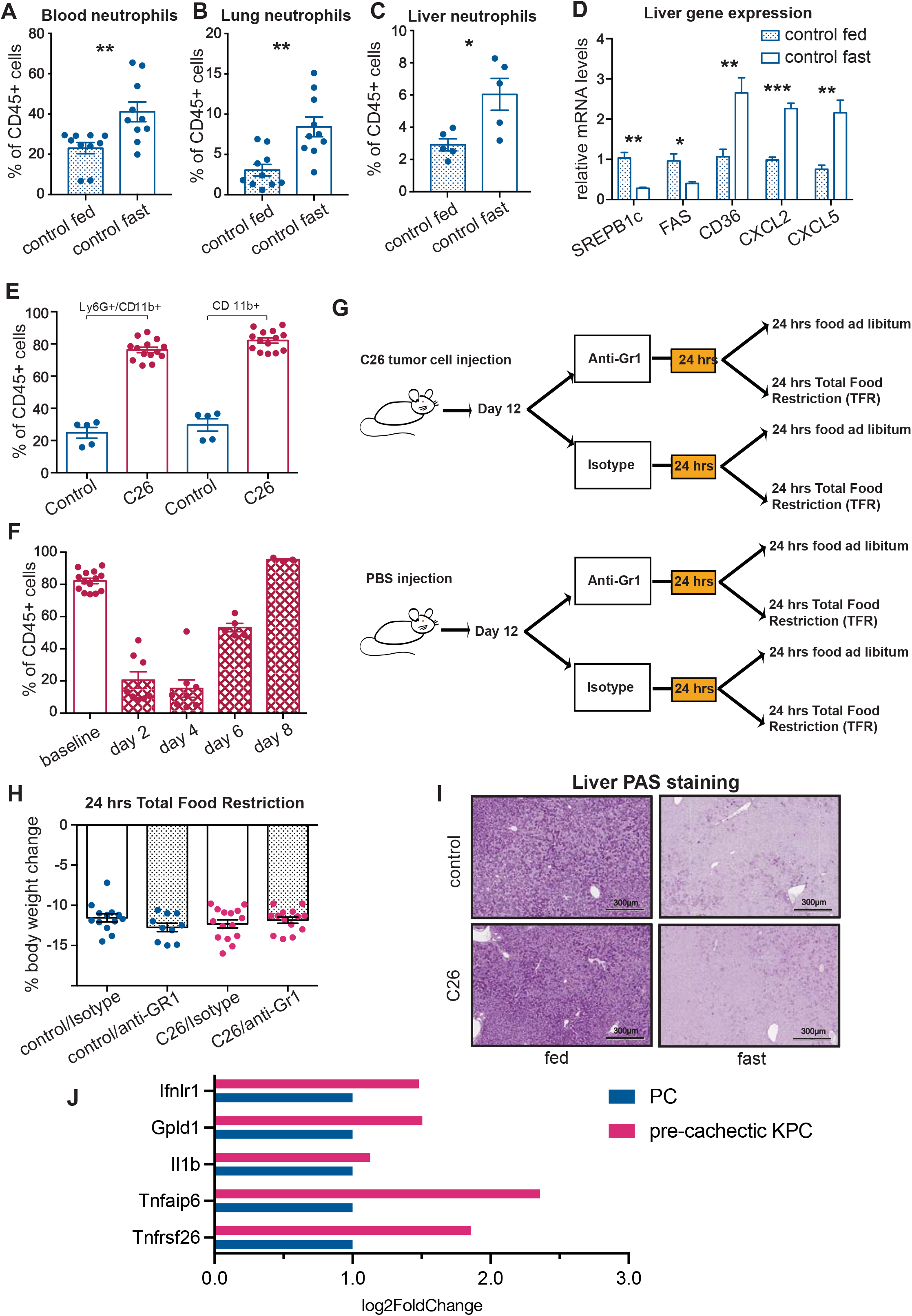
Immune characterization of the C26 mouse model of cachexia. (A-C) Quantification of neutrophils (displayed as % of all CD45+ cells) by flow cytometry in the blood (A), lung (B), and liver (C) of control mice fed and after 24 hours of total food restriction (TFR); (D) Gene expression levels of metabolic genes and chemoattractant ligands in the liver of mice in Fig EV3C-E; (E) Quantification by flow cytometry of Ly6G+/CD11b+ and CD11b+ cells (displayed as % out of all CD45+ cells) in the blood of C26 mice and littermates; (F) Quantification by flow cytometry of CD11b+ (displayed as % out of all CD45+ cells) in the blood of C26 mice at baseline and every other day after starting treatment with anti-Gr1; (G) Schematic representation of the TFR and neutrophil depletion experiments; (H) Percentage body weight changes after TFR in C26 mice and controls treated with anti-Gr1 or isotype; (I) Periodic acid-Schiff (PAS) staining of glycogen in the liver of fed or fasted pre-cachectic C26 mice and controls; (J) Gene expression analysis of selected lipolytic genes by RNA sequencing as in Fig 3A: IFNLR1 (Interferon Lambda Receptor 1). Phosphatidylinositol-glycan-specific phospholipase D (GPLD1), a hydrolyzing enzyme that breaks the linkage of proteins to the cell membrane thereby releasing protein molecules and modifying glycemic indexes. Interleukin 1 Beta (IL1β), a lipolytic cytokine that mediates the inflammatory response. Tumor necrosis factor-inducible gene 6 protein (TNFAIP6) (related to lipolytic cytokine TNF), gene associated with inflammation and cell migration. Tumor necrosis factor receptor superfamily member 26 (TNFAIP26), involved in inflammation and cell survival.

